# Ketamine reverses stress-induced hypersensitivity to sunk costs

**DOI:** 10.1101/2024.05.12.593597

**Authors:** Romain Durand-de Cuttoli, Brian M. Sweis

## Abstract

How mood interacts with information processing in the brain is thought to mediate the maladaptive behaviors observed in depressed individuals. However, the neural mechanisms underlying impairments in emotion-cognition interactions are poorly understood. This includes influencing the balance between how past-sensitive vs. future-looking one is during decision-making. Recent insights from the field of neuroeconomics offer novel approaches to study changes in such valuation processes in a manner that is biologically tractable and readily translatable across species. We recently discovered that rodents are sensitive to “sunk costs” – a feature of higher cognition previously thought to be unique to humans. The sunk costs bias describes the phenomenon in which an individual overvalues and escalates commitment to continuing an ongoing endeavor, even if suboptimal, as a function of irrecoverable past (sunk) losses – information that, according to classic economic theory, should be ignored. In the present study, mice were exposed to chronic social defeat stress paradigm, a well-established animal model used for the study of depression. Mice were then tested on our longitudinal neuroeconomic foraging task, Restaurant Row. We found mice exposed to this severe stressor displayed an increased sensitivity to sunk costs, without altering overall willingness to wait. Mice were then randomly assigned to receive a single intraperitoneal injection of either saline or ketamine (20 mg/kg). We discovered that stress-induced hypersensitivity to sunk costs was renormalized following a single dose of ketamine. Interestingly, in non-defeated mice, ketamine treatment completely abolished sunk cost sensitivity, causing mice to no longer value irrecoverable losses during re-evaluation decisions who instead based choices solely on the future investment required to obtain a goal. These findings suggest that the antidepressant effects of ketamine may be mediated in part through changes in the processing of past-sensitive information during on-going decision-making, reducing its weight as a potential source of cognitive dissonance that could modulate behavior and instead promoting more future-thinking behavior.

## Introduction

At a fundamental level, emotional states are capable of augmenting the way information is processed in the brain, influencing how actions are selected.^1^ This is generally thought to serve adaptive purposes, for example with hunger state or threat perception biasing action-selection processes that promote safety and survival. However, in the context of psychiatric illnesses, emotion-cognition interactions can become dysfunctional and maladaptive.^2^ Depression is a severely debilitating mental illness characterized by persistent emotional dysregulation, impacting an estimated 17.3 million adults in the United States in 2017 alone, 7.1% of the adult population.^3,4^ Although changes in mood is the most striking clinical feature of this disorder, depression can substantially impair quality of life by affecting attention, motivation, and cognition that in turn drive maladaptive changes in behavior and can perpetuate this illness.^3^ Patients with depression can have impaired decision-making.^5^ However, little is known about the relationship between mood and decision-making information processing in the brain of depressed individuals.

It is intuitive to appreciate how decisions and their outcomes can augment mood (e.g., related to surprise, disappointment, regret, etc.), however it is less apparent for individuals to appreciate how their choices are conversely being influenced by mood.^6^ Emotions can drive individuals to make decisions they typically would not. For instance, emotional state can augment the perceived value of wins differently from losses.^7^ Psychologists and behavioral economists have long observed that people are more sensitive to losses than they subjectively value objectively equivalent gains.^8^ This cognitive phenomenon has been well-explained by a concept known as prospect theory, which describes how individuals are generally more loss averse in their decision-making characteristics.^7,8^ This is often a result of an accumulated history of experiences in which an individual has suffered losses in the past, each episode of which might carry strong emotional salience and, in turn, drive individuals to use that information to guide future behavior. According to normative models of decision-making, individuals should only evaluate the expected future utility of a chosen action when deliberating amongst options.^9^ However, individuals are rarely neutral about choice history and are particularly sensitive to past losses when making future decisions.^10^

A cognitive phenomenon known as the sunk cost bias captures a concrete example of a valuation process in which information about the past that, economically speaking, should bear no weight in a choice can in fact powerfully augment decision-making.^10,11^ The sunk cost bias describes the cognitive phenomenon in which an individual escalates the commitment of an ongoing endeavor based on investments (e.g., of time, money, effort, etc.) already made toward a goal independent of future prospects.^11^ That is, individuals overvalue an option not solely based on the effort required to obtain a goal but rather the effort already expended, which theoretically should be ignored as it is an irrecoverable (or sunk) cost. Thus, an economically “rational” decision made between two competing options should only consider the predicted value of those two alternative futures, ignoring the unchangeable past since that factor will remain the same in either case. Yet, individuals often honor sunk costs for reasons that remain unclear but are thought to be linked to minimizing a source of cognitive dissonance.^12^ Focusing on prior losses can generate negative affect that can be somewhat ameliorated if an individual can justify prior decisions by not abandoning an ongoing endeavor, which may be otherwise perceived as a failure on the part of the decision-maker.^13^ While humans are sensitive to this information to variable degrees, this valuation process may be pathological in depression.^14^ Propensity to perceive negative affect, heightened in depressed individuals, has been associated with increased sensitivity to sunk costs.^15^ Leahy’s sunk cost theory of depression suggests that because of this exaggerated process patients form a barrier to change.^14,16^ This may be related to what is sometimes referred to as inaction inertia and regret avoidance in the face of uncertainty that may be exacerbated in depressed individuals.^6,17,18^ For patients, these behaviors may be rooted in a failure to disengage in the face of perceived heavy costs associated with the risk of failure. Thus, Leahy’s sunk cost theory of depression is thought to be a reason why depressed patients remain locked into repeated patterns of self-defeat.^14,16^

Psychological theories of depression have been extremely useful for guiding cognitive behavioral therapy and offering a framework for which to investigate the neural correlates of emotional dysregulation.^19^ However, much of the discoveries in the neurobiology of depression stem from non-human animal research that, while instrumental in aiding the testing of psychopharmacology and newer neuromodulation treatment options, their ability to approximate the complexity of human features of depression is often limited and thus a fundamental roadblock to gaining translational insights into brain-behavior relationships that are clinically relevant.^20–25^ Recent discoveries in the field of neuroeconomics have begun to push the boundaries on what can be investigated in animal research, offering novel approaches using clever tasks to extract hidden features of behavior in order to study interactions between mood and decision-making in a manner that is biologically tractable and readily translatable across species.^26–33^ The sunk cost bias has been extensively studied in the human psychology literature but has been repeatedly ascertained that this phenomenon is unique to humans.^34,35^ Using a neuroeconomic approach, it was recently discovered for the first time evidence that rodents are capable of demonstrating sensitivity to sunk costs in a manner that is directly matched to human behavior using a novel set of neuroeconomic decision-making tasks recently developed for use across species.^36^ These findings suggest there is a conserved evolutionary history to such cognitive biases that arise from similar neural systems across species, opening a door to a novel line of scientific inquiry in translational behavioral neuroscience and offering a new approach to characterize the neural systems involved in emotion-cognition interactions.

To date, this neuroeconomic approach to study complex decision-making in rodents has only recently been applied in the context of depression.^37–39^ Furthermore, the biological and psychological mechanisms through which antidepressant treatments, including newer rapid-acting drugs like ketamine, exert their effects are not well understood.^33,40–42^ Here, we tested the effects of stress using a well-established animal model of depression (the chronic social defeat stress paradigm) on mice tested in a novel neuroeconomic decision-making paradigm (the Restaurant Row task) in which mice forage for food rewards of varying costs while on a limited budget.^36,43^ We found that exposure to a psychosocial stressor produced a robust increase in sensitivity to sunk costs without affecting gross changes in locomotion, motivation, body weight, or feeding behaviors. We also found that following a single dose of ketamine, this hypersensitivity to sunk costs was renormalized in stressed mice and completely abolished in non-stressed mice. Our findings demonstrate a novel psychological mechanism of how a relatively new antidepressant agent such as ketamine may selectively access complex emotion-decision interactions in order to exert its therapeutic effects, shedding light on the psychological computations that may be affected by depression and involved in its recovery. Together, these data demonstrate how utilizing neuroeconomic approaches to decision-making can enhance the richness of translational, behavioral endpoints in animal studies in efforts to better approximate human cognition.

## Methods

### Animals

In this study, 60 male C57BL/6J were purchased from Jackson Laboratories at 11 weeks of age. 44 of these animals were put through the chronic social defeat stress protocol (CSDS, 10 days) described below while 16 mice served as non-stressed (non-defeated) controls. 22 of the 44 defeated mice and 10 of the 16 non-defeated mice were promoted to go on to neuroeconomic decision-making testing (60 days) selected based on matched bodyweights, totaling 32 mice that advanced to the neuroeconomic decision-making testing described below. Half of each of the defeated and non-defeated groups were then either injected with ketamine or saline outside of behavioral testing hours before being tested for an additional 5 days on the neuroeconomics decision-making task. Mice were then sacrificed for tissue harvesting. Experimental timeline is depicted in **Figure 1A**.

**Figure 1.**
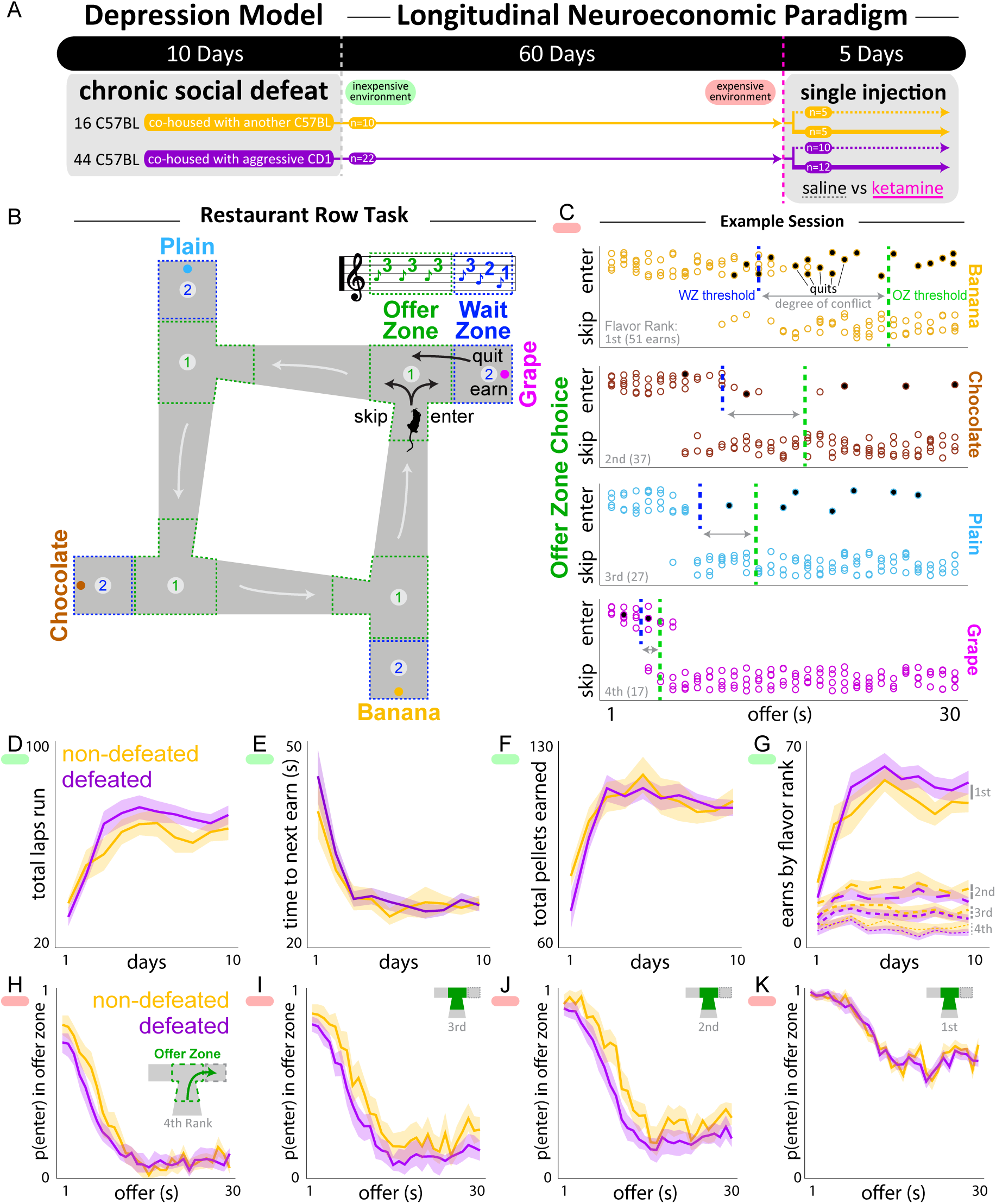
Mice exposed to the chronic social defeat stress (CSDS) model of depression are capable of learning the neuroeconomic foraging task, Restaurant Row. (A) Experimental timeline. Mice underwent 10 days of the chronic social defeat stress model of depression being paired either with an aggressive CD-1 mouse (defeated) or another C57BL/6J mouse (non-defeated controls). A subset of mice then underwent 60 days of training on the Restaurant Row task that escalated from an inexpensive reward rich environment early on in training (green epoch, offers ≤ 5s during the first 10 days) to an expensive reward scarce environment late in training (red epoch, offers ranging from 1s to 30s). Subsets of mice were then injected with a single dose of either ketamine or saline and then were tested for another 5 days on the task before being sacrificed. (B) Task schematic. Food-restricted rodents were trained on a maze encountering serial offers for flavored rewards in four “restaurants.” Each restaurant contained a separate offer zone and wait zone. Tones sounded in the offer zone; fixed tone pitch indicated delay rodents would have to wait in the wait zone (up to 30s, random on offer zone entry). Tone pitch descended in the wait zone during delay “countdown.” Rodents could quit the wait zone for the next restaurant during the countdown, terminating the trial. (C) Data from an example 60min session from a well-trained mouse in an expensive reward scarce environment (red epoch). Dots represent individual trial choices to skip or enter in the offer zone of each restaurant as a function of cued offer. Enter trials that end as quits are filled in black. Flavor preference is ranked by summing total earns in each restaurant at the end of each day (gray text, 1^st^ is most preferred). Dashed lines represent the indifference points fit by a sigmoid function of choices in either the offer zone (green line) or wait zone (blue line). Gray arrows highlight the degree of discrepancy between willingness to enter (offer zone [OZ] threshold) and willingness to wait (wait zone [WZ] threshold). (D-H) Basic behavior in an inexpensive reward rich environment (green epoch). (D) Average total laps run in the correct direction. (E) Average time elapsed between successive pellets earned. (F) Average total number of pellets earned. (G) Average body weight measured immediately before (dashed) and after (solid) testing. (H) Average number of pellets earned in each restaurant ranked by flavor preference (1^st^, most preferred, solid). (I-L) Offer zone deliberation behavior in an expensive reward scarce environment (red epoch). Mice are able to make economically advantageous decisions to enter affordable offers cued by tone pitch plotting probability of entering as a function of cost interacting with flavor preferences (I, 4^th^, least preferred – L, 1^st^, most preferred). Shaded lines represent mean ±1SEM.

### Husbandry

Mice were group housed until they began this experiment. During the CSDS protocol mice were co-housed with one other mouse for 10 days before being individually housed for the remainder of this experiment. Mice had access to water ad libitum during this entire experiment. Prior to and during the CSDS paradigm, mice had access to regular chow ad libitum. Following the CSDS paradigm, mice were then food restricted to 85% of their free-feeding body weight and switched to a flavored pellet diet (BioServe products). These 20mg pellets were used in the neuroeconomic decision-making paradigm as operant food rewards automatically delivered through a pellet dispenser and served as these animals’ sole source of food. These pellets are full-nutrition and vary only in flavor (either chocolate, banana, grape, or plain). Mice spent days following the end of the CSDS paradigm acclimating to individual housing and pellet trained in their home cage introduced to a single daily serving (3g) of the flavored pellets before beginning training on the neuroeconomic decision-making task. Mice were weighed daily during the CSDS paradigm and twice daily during the neuroeconomic decision-making task (before and after testing). All mice were on a regular 12-hour light, 12-hour dark cycle from 7am to 7pm housed in the temperature – and humidity-controlled Mount Sinai Annenberg Animal Care Facility. Experiments were approved by the Mount Sinai Institutional Animal Care and Use Committee (IACUC; protocol number LA12-00051) and adhered to the National Institutes of Health (NIH) guidelines.

### Depression Model: The Chronic Social Defeat Stress (CSDS) Paradigm

Mice underwent the CSDS paradigm, a well-established animal model of psychosocial stress that is capable of inducing a depressive-like phenotype in mice.^43^ This paradigm involved co-housing the C57BL/6J mouse with a 4-6-month-old retired CD-1 breeder mouse, an outbred strain that tend to exhibit overly aggressive behavior toward other mice. In this protocol, the C57BL/6J mouse became subordinates in this social hierarchy. Both mice were allowed to interact for 5 minutes during which the C57BL/6J mouse was attacked by the CD-1 mouse. So as to prevent bodily injury, mice were then separated and for the remainder of the day were co-housed in the same cage but separated by a mesh divider so the mice no longer had direct physical contact but continued to have visual, olfactory, and auditory contact. This living environment placed the C57BL/6J mouse in a constant state of psychosocial stress. This procedure was repeated for 10 consecutive days with 10 different CD-1 mice (**Figure 1A**). As a control to this stressor, non-stressed (non-defeated) C57BL/6J mice were handled equally without exposure to CD-1 mice but were exposed to other, domiciled C57BL/6J mice of the same size and age. By the end of this protocol, mice are typically assayed on a number of simple behavioral tests (sucrose preference test, elevated plus maze, social interaction test) that collectively capture depression-related phenotypes (anhedonia, anxiety, social avoidance, respectively).^43^ However, the goal in this study, rather than to assay mice on these simple behavioral protocols, was to test defeated and non-defeated mice on the novel neuroeconomic decision-making paradigm, the Restaurant Row task, in order to characterize complex choice behavior.

### Neuroeconomic Decision-Making Paradigm: The Restaurant Row Task

In the Restaurant Row task, mice were trained to forage in a square maze for food rewards of varying cost (delays ranging from 1 to 30s) and subjective value (unique flavors tied to four separate and uniquely spatially cued locations, or restaurants, located in each corner of the maze) while on a daily limited time budget (60 minutes). Each restaurant consisted of two separate decision zones, (1) an offer zone and (2) a wait zone (**Figure 1B**). Upon entry into a restaurant’s initial offer zone, a tone sounded whose pitch indicated the delay mice would have to wait in order to earn food if they chose to enter the wait zone (higher tone pitch equates to a longer delay randomly selected upon offer zone entry; pitch identities were shared across restaurants). If a mouse chose to enter the wait zone, a countdown began during which tones descend stepwise every second either until the reward was earned or the mouse decided to quit and leave the wait zone. There is no penalty to quitting other than the offer was rescinded and the mouse must advance to the next restaurant. Thus, a trial was terminated if a mouse made (1) a skip decision in the offer zone, (2) a quit decision in the wait zone, or (3) earned a reward, after which the mouse must progress to the next restaurant in a serial order (**Figure 1C**). Importantly, rewards earned on this task served as the only source of food (full nutrition flavored BioServe pellets), making this task closed-economic in nature with time as a limited commodity. This means decisions made on this task were interdependent both across trials as well as across days. This also means that any time spent engaged in a given behavior was at the expense of spending from the time budget engaged in other behaviors or exploring alternative options, a concept known as opportunity cost in the foraging literature.^44,45^ Thus, how value is calculated can be operationalized in many forms depending on available actions to choose from, choice history, and current economic situation. Different animals preferred the unique flavors differently. Flavors were used to modulate subjective value determined by revealed preferences without assuming reward value (as opposed to varying pellet number in each restaurant that comes with increased handling time per pellet). Flavor preferences were determined by summing the total pellets earned in each restaurant and ranking flavors from most preferred (1^st^) to least preferred (4^th^) each day (**Figure 1C**). Flavor preferences developed early in training and were stable across days, consistent with previously published reports.^46,47^

### Ketamine Treatment

After day 60 of testing, animals were randomly assigned to receive either saline or ketamine, counterbalancing groups across as many behavioral parameters as possible. After 60 days of training mice were either injected with saline (0.9% NaCl) or 20mg/kg ketamine intraperitoneally. Restaurant Row testing always took place during the day during the animals’ light phase. Only on one special day when injections were to be administered, injections took place in the dark phase in the evening after Restaurant Row testing for that day was completed. All injections were volume corrected measuring mouse body weights immediately before injections. Restaurant Row testing resumed the next day for 5 additional days. Overall, our goal was to measure how decision processes were affected following a single dose of ketamine versus saline after the acute effects of the drug have worn off and ketamine was no longer in the animals’ systems (the half-life of ketamine is 45 minutes).^48^

## Results

All mice were able to learn the general structure of the Restaurant Row task. That is, in the first 10 days of training, all mice were able to learn to run laps in the correct direction (a full counterclockwise loop around the maze trigging one offer in each restaurant), quickly increasing to a stable number of laps run over the first few days (approximately 80 laps or 320 total trials [individual offers]) consistent with previously published reports and with no significant differences between defeated and non-defeated mice (main effect across days: F_31,9_=80.54, p<0.0001, interaction day x CSDS: F_9,1_=1.92, p=0.17, **Figure 1D**).^47^ Early on in training, when costs were relatively inexpensive (less than or equal to 5s delays at most) mice were able to quickly earn pellets at a stable reinforcement rate of approximately 30s in between subsequent earns, with no significant differences between defeated and non-defeated mice (main effect across days: F_31,9_=58.24, p<0.0001, interaction day x CSDS: F_9,1_=1.44, p=0.23, **Figure 1E**). In doing so, mice were able to achieve a stable number of pellets earned and body weight gained within a few days leveling off at approximately 100 pellets (or 2g) earned on the task every day (enough food to sustain healthy nutrition based on previously published reports), with no significant differences between defeated and non-defeated mice (<u>total pellets earned:</u> main effect across days: F_31,9_=23.97, p<0.0001, interaction day x CSDS: F_9,1_=1.62, p=0.20; <u>post-task body weight:</u> main effect across days: F_31,9_=12.94, p<0.001, interaction day x CSDS: F_9,1_=1.49, p=0.223, **Figure 1F-G**).^47^ Within a few days of starting the Restaurant Row task, mice revealed flavor preferences that remained stable throughout testing, with mice strongly preferring one flavor over the others earning approximately 60 pellets in their most preferred restaurant and maintaining ordinal rankings of subjective value across flavors, with no significant differences between defeated and non-defeated mice (two-way interaction day x flavor ranking: F_9,3_=23.22, p<0.0001; three-way interaction day x flavor ranking x CSDS: F_3,1_=1.27, p=0.28, **Figure 1H**). With extended training, mice advanced to a more reward-scarce environment in which offers ranged anywhere from 1s to 30s randomly selected upon entry into each restaurant’s offer zone. After prolonged training, mice were capable of learning how to successfully discriminate among tone pitches upon entry into a restaurant’s offer zone, reliably entering low-delay offers and skipping long-delay offers as a continuous function of delay as well as a function of flavor preference with no significant differences between defeated and non-defeated mice (two-way interaction offer x flavor ranking: F_29,3_=13.17, p<0.0001; three-way interaction offer x flavor ranking x CSDS: F_3,1_=0.81, p=0.49, **Figure 1I-L**). The fact that mice treated the tone pitches differently across restaurants indicates that mice are able to engage in a flexible goal-directed valuation process integrating offer cost (auditory cue indicating the delay of a given trial) and subjective value (learned spatial context indicating the flavor of a given trial). Taken together, these data demonstrate that chronic social defeat stress did not have any significant impact in impairing how mice grossly perform on the Restaurant Row task. Importantly, these data demonstrate that in a naturalistic foraging paradigm, a mouse model of depression that has been previously able to produce changes in phenotypes of anhedonia, amotivation, feeding behavior, and locomotor activity in mice failed to disrupt gross signs of these processes on the Restaurant Row task and that defeated mice, in fact are able to engage in goal-directed behaviors with locomotor, feeding, hedonic valuation, and planning capabilities seemingly intact.^43^

To operationalize decision-making behaviors on this task related to sunk costs, we specifically examined choices in the wait zone. Choosing between offers in the offer zone (making enter versus skip decisions as a function of cost and flavor) are thought to access fundamentally distinct decision-making processes from the choices made in the wait zone (deciding to remain patient and earning a reward (inaction) versus opting out of an ongoing investment and quitting (action).^45,49–51^ These wait zone decisions have been previously linked to more affective components of decision-making information processing in which animals must decide to abandon an ongoing investment and backtrack.^36,46,47^ The sunk cost bias describes how investments made to an ongoing endeavor are capable of escalating the commitment of continuing with that endeavor.^10^ This process can be directly captured in the decision dilemma mice face in the wait zone. Indeed, using the analysis described below, it was previously demonstrated that healthy mice, rats, and humans show sensitivity to sunk costs on the Restaurant Row task and the human equivalent task.^36^

To measure the sunk cost effect, we first calculated the probability mice earn rewards in the wait zone as a function of offer length immediately upon entry into the wait zone (**Figure 2A**). As expected, mice overall have an increased probability of earning a reward (less likely to quit) the shorter the starting delay (**Figure 2A**). Here, the probability of earning versus quitting is calculated at the onset of tone countdown beginning in the wait zone. Next, we then analyzed a subset of trials defined both by the initial starting offer accepted as well as the amount of time already waited in the wait zone. For example, if a mouse entered a 10s offer, and waited 5s of the countdown thus far (5s sunk), with 5s remaining in the countdown required to earn the reward, the probability of earning versus quitting can be calculated at this specific moment in time (analysis point, **Figure 2A**). By comparing this scenario to another trial in which the mouse had entered a 15s offer and waited 10s (10s sunk, with 5s remaining in the countdown) or had just entered a 5s offer and not yet waited at all (0s sunk), the effect of time already waited on continuing to invest in an ongoing endeavor can be observed independent of and separate from temporal distance to the goal, or future effort required (in this example, three scenarios all matched to 5s remaining in the countdown). This analysis can be repeated as a continuous function of time remaining in the countdown (future effort required) and time already waited (sunk costs) by examining every combination of starting offers and varying the analysis point.

**Figure 2.**
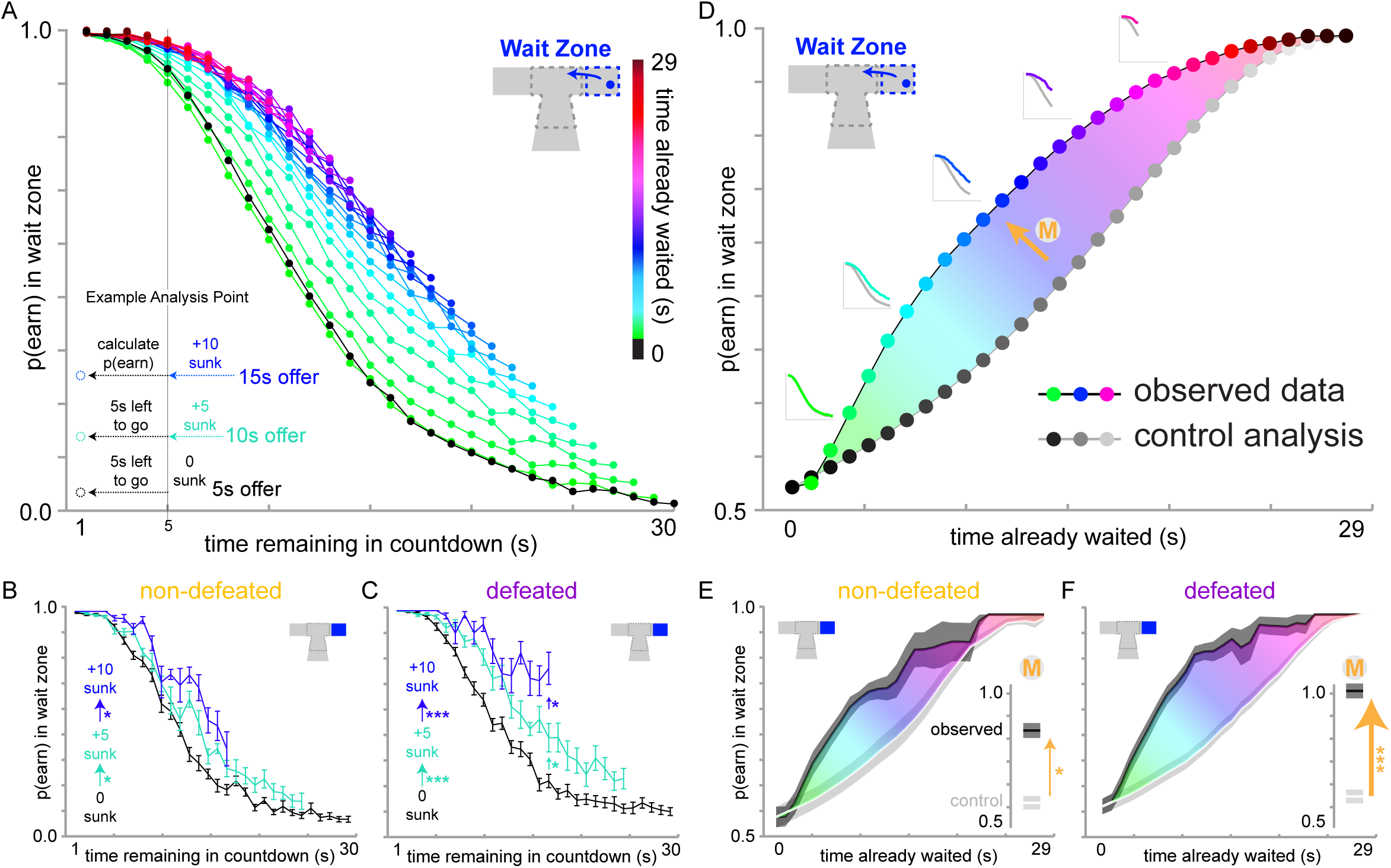
Chronic social defeat stress (CSDS) enhances sensitivity to sunk costs during change-of-mind decisions in the wait zone. (A) Wait zone sunk cost analysis demonstrated on data from all mice pooled together. X axis reflects the cued time left remaining in the countdown. Y axis represents the calculated probability of completing the countdown. Inset depicts a visual explanation of an example analysis point comparing three sunk cost conditions (0s sunk, black; 5s sunk, teal; 10s sunk, blue) all matched with the same amount of time remaining in the countdown (5s left to go, thus the initial starting offer that was entered in these example conditions were 5s black, 10s teal, and 15s blue, respectively). Vertical deviation from black 0s sunk curve as a function of time already waited (color) reflects the sunk cost-driven inflation of the value of continuing to wait. (B-C) Grouped data from three example sunk cost conditions in (B) non-defeated and (C) defeated mice. (D-F) Second-order dimension-reduced analysis collapsing across time remaining in countdown for each sunk cost condition and instead only displaying p(earn) as a function of time already waited. Insets show which curves from panel (A) are collapsed horizontally into a single data point in (D) at every given X value. Control analysis represents collapsed data from the black 0s sunk curve in (A) leaving out the right most datapoint re-iteratively to match the number of datapoints collapsed into each sunk cost color (observed data). The magnitude of the sunk cost effect (“M”, gold arrow) is the orthogonal distance between the observed and control analysis curves. (E-F) Grouped data across the continuum of sunk cost conditions in (E) non-defeated and (F) defeated mice. Insets summarize magnitude scores. Error bars and shaded lines represent mean ±1SEM. *p<0.01, ***p<0.0001.

Here, we replicated previously published findings demonstrating that mice are sensitive to sunk costs (**Figure 2A**).^36^ A critical tenant of the sunk cost bias is not only that prior investments can inflate the value of continuing to invest in an ongoing endeavor but that this effect is stronger the more prior losses that have been incurred.^10^ This facet of the sunk cost bias is captured in the wait zone as a function of time already waited further increasing the probability of earning a reward in the wait zone. Interestingly, we found that defeated mice displayed an increased sensitivity to sunk costs compared to non-defeated mice (<u>defeated mice:</u> interaction time remaining in countdown x time already invested at 0s, 5s, and 10s sunk conditions: F_29,2_=19.59, p<0.0001; <u>non-defeated mice:</u> interaction time remaining in countdown x time already invested at 0s, 5s, and 10s sunk conditions: F_29,2_=5.59, p<0.01, **Figure 2B-C**) independent of temporal distance to the goal. To integrate and summarize the sunk cost effect across all permutations of trials, we carried out a subsequent analysis that collapses the probability of earning across time remaining in the countdown reducing the dimensionality of this analysis to focus instead specifically on sunk costs (**Figure 2D**). As a control analysis, the 0s cost curve can be re-iteratively collapsed on itself across time remaining in the countdown, leaving out the right most data point to match datasets for each respective sunk cost condition that is collapsed across time remaining in the countdown. Thus, in this second-order analysis, the magnitude of the sunk cost effect can be directly measured as area between the dimension-reduced sunk cost curve and the control analysis curve (**Figure 2D**). In this second-order analysis, we found that as a continuous function of sunk costs, defeated mice display an increased sensitivity to sunk costs (three-way interaction time already invested x analysis curve x CSDS: F_3,1_=5.71, p<0.05, **Figure 2E-F**).

Following 60 days of testing on the Restaurant Row task, mice were injected with a single dose of ketamine or saline and subsequently retested on Restaurant Row (**Figure 3**). We found that the magnitude of sunk costs was reduced in defeated mice that received an injection of ketamine (F_28,1_=18.61, p<0.01, **Figure 3D**) while defeated mice who were treated with saline continued to display hypersensitivity to sunk costs (F_28,1_=100.97, p<0.0001, **Figure 3B**). Interestingly, non-defeated mice who were treated with ketamine too displayed a reduction in the magnitude of sunk costs such that these mice were no longer sensitive to sunk costs (<u>ketamine</u>: F_28,1_=0.07, p=0.80, **Figure 3C**; <u>saline</u>: F_28,1_=22.98, p<0.01, **Figure 3A**). These data suggest that a single dose of ketamine is capable of augmenting a higher-order valuation process such as propensity to honor sunk costs in a manner that renormalizes this cognitive phenomenon in mice with a history of psychosocial stress exposure while completely abolishing this valuation algorithm in healthy mice with no psychosocial stress history.

**Figure 3.**
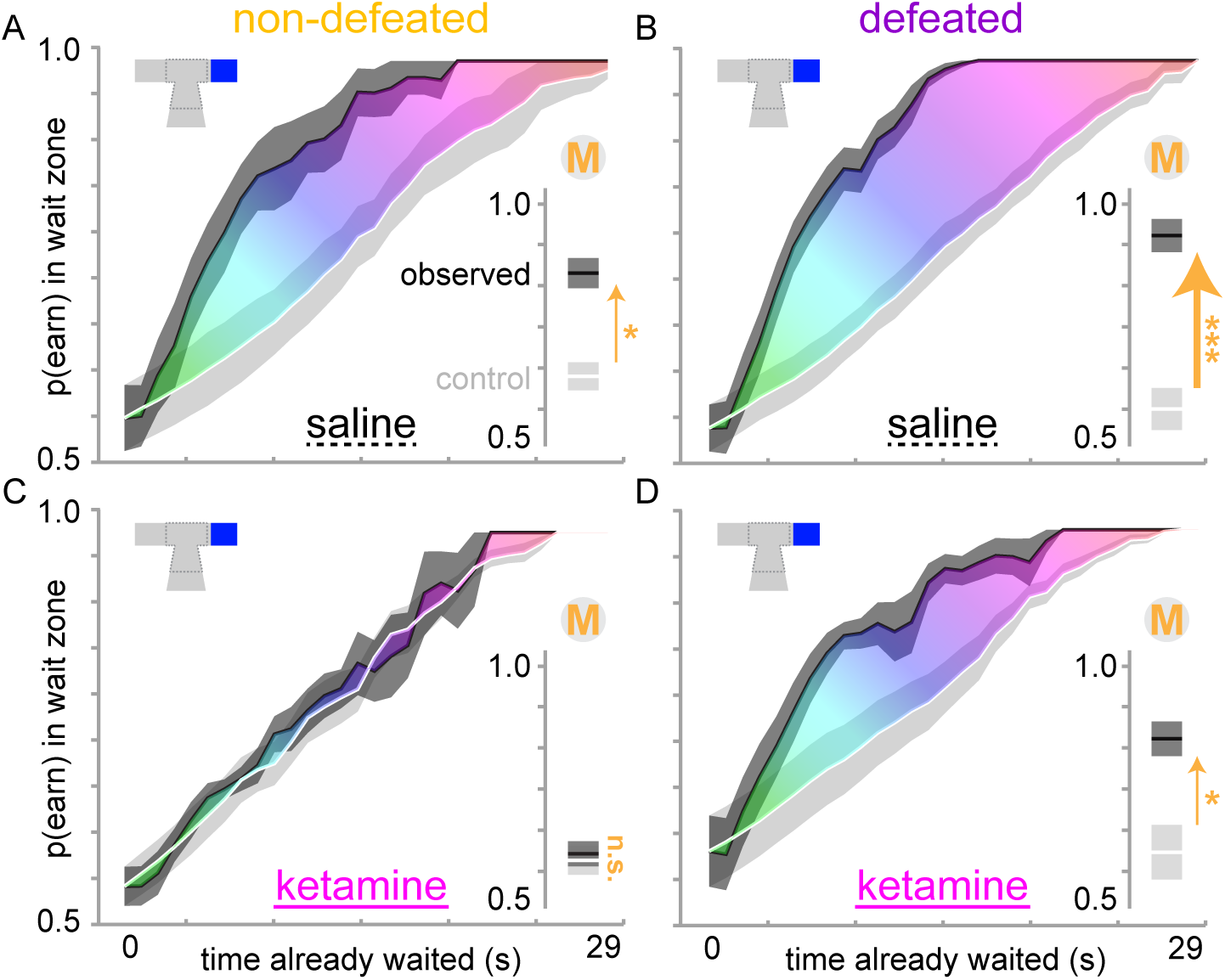
A single dose of ketamine renormalizes hypersensitivity to sunk costs in defeated mice while abolishing sunk costs in non-defeated mice. Same second-order dimension-reduced analyses as depicted in Figure 2D-F 24 hours following one-time injections with either saline (A-B) or ketamine (C-D) in either non-defeated (A,C) or defeated (B,D) mice. Insets summarize the magnitude of the sunk cost effect (“M”, gold arrow, described in Figure 2D). Shaded lines represent mean ±1SEM. Not significant, n.s., p>0.05. *p<0.01, ***p<0.0001.

## Discussion

In this study, we set out to investigate the effects of psychosocial stress on higher-order emotion-cognition interactions. Specifically, we tested mice following exposure to stress in the CSDS paradigm, a well-established animal model of depression, on the neuroeconomic Restaurant Row decision-making task.^43^ The Restaurant Row task is a naturalistic foraging paradigm developed for use in rodents and translated across species for use in humans that has recently demonstrated promise in being able to access complex decision-making processes measured via subtle behavior.^36,47,49,50,52–55^ Sophisticated analyses of behavior on this task have been able to extract hidden features of information processing such that cognitive phenomena like the sunk cost bias can be directly measured.^36^ The sunk cost bias describes the phenomenon in which individuals pay special attention to irrecoverable past losses that should otherwise be ignored in their decision-making pursuits, driving the escalation of commitment to ongoing endeavors, even those that may be detrimental to own’s own interests.^10^ This decision-making phenomenon has recently been discovered in non-human species and offers a way to study complex emotion-cognition interactions in a laboratory setting.^36^ Using this neuroeconomic approach in mice, we discovered that a history of psychosocial stress produced a hypersensitivity to sunk costs and that this phenotype could be reversed by a single dose of ketamine.

The CSDS model of depression has been widely used by many laboratories investigating the effects of stress on the body, brain, and behavior.^43^ Unlike other stress protocols commonly used in animal studies (e.g., physical restraint stress, variable stress, early life maternal separation stress, chemical stress, etc.), this model captures elements of social subordination which carries ethological relevance and face validity in its ability to approximate stress-inducing psychosocial relationships and recapitulate lasting symptoms of human depression in mice, including but not limited to alterations in motivational state, anhedonia, anxiety-like behaviors, despair-related behaviors, social avoidance, drug-use lability, circadian rhythm shifts, and other metabolic dysfunction seen in clinically depressed populations.^3,43^ Behavioral deficits induced by this stress protocol have been shown to be reversed by chronic but not acute treatment with antidepressants across a number of different drug classes, particularly monoamine-based psychopharmacologic therapies including selective serotonin reuptake inhibitors, serotonin norepinephrine reuptake inhibitors, monoamine oxidase inhibitors, and tricyclic antidepressants.^56,57^ This model has afforded the ability to explore neurobiological changes induced by psychosocial stress and augmented by antidepressant treatments in an experimental laboratory setting with the cutting-edge tools developed for mice available today in modern neuroscience.^20,43,57^

Discoveries that have stemmed from such animal models of depression cover wide areas of research spanning from cellular and molecular to circuit-level and systems neuroscience.^20,25,57^ Current theories of depression as well as hypotheses of the therapeutic mechanisms of antidepressants available today that have stemmed from this line of research are largely based on dysregulation of the hypothalamic-pituitary-adrenal axis as well as several regions involved in the brain’s reward and limbic systems affecting biological processes in the endocrine, electrophysiological, genetic, and more recently discovered, epigenetic domains of function.^22,58–61^ For example, many studies over the past two decades have investigated the impact of a repeated social stress on the neural pathway connecting the ventral tegmental area (VTA) and the nucleus accumbens (NAc) regions of the brain.^20^ This VTA-NAc mesolimbic dopaminergic pathway has been shown to play a key role in the integration of both rewards and emotions.^20^ Major findings in the cellular and molecular adaptations in this pathway in response to stress include (but are not limited to): identifying the role of brain-derived neurotrophic factor (BDNF) as a key regulator of the genetic adaptations following CSDS and its function in driving an increase in the firing patterns of dopaminergic neurons in the VTA associated with CSDS, the role of the transcription factor DeltaFosB in protecting against depressive-live phenotypes, and the role of RAS-related C3 botulinum toxin substrate 1 in the NAc and its epigenetic regulation in promoting increased susceptibility to stress.^43,62–67^ Recently, individual differences in the peripheral immune system have been shown to serve as major modulators of susceptibility to stress.^68^ Stress-induced neurovascular disruption of the blood-brain barrier in the NAc has been recently shown to serve as a potential mechanism linking peripheral stress responses to impacts in the central nervous system.^69–72^ Despite these major advances, which have made promising strides in beginning to inform novel targets for drug development, new strategies for neuromodulation intervention, individualized approaches to pharmacological treatment paths, and even expanding our understanding of the heritability of depression, animal models of depression still struggle with bridging the translational gap toward understanding the complexities of human emotion and how neurobiological changes underlie psychological features of depression. The Research Domain Criteria Initiative from the National Institute of Mental Health has been calling for an overhaul in the way the field approaches the study of functional domains of mental health that span across multiple levels of analysis.^4^ Part of what has slowed progress in making significant steps toward this end is the reliance on simple behavioral paradigms in animal models which have changed very little relative to the recent explosion of innovative tools available in neuroscience today.

Neuroeconomics has recently demonstrated how novel approaches to non-human behavioral studies can move beyond simple tests of value and in turn reveal more about complex emotion-cognition interactions.^28,49^ In this study, we trained mice on the Restaurant Row task that has been previously validated across species to healthy humans on a matched version of the task.^36,53^ We found that mice exposed to CSDS displayed no gross impairments in locomotion or motivated feeding behaviors and were capable of learning the structure of the Restaurant Row task. At first glance, this may appear at odds with existing literature showing how stress can promote anhedonic phenotypes in mice measured using sucrose preference tests.^43^ Unlike simple tasks such as the sucrose preference test, the Restaurant Row task may be accessing fundamentally distinct or more subtle and more narrowly defined valuation algorithms used in reward pursuit than those observed by simply measuring sucrose consumption in the home cage. That is, the Restaurant Row task tests mice in a more naturalistic foraging setting with a more dynamic and engaging environment pitting multiple competing decision processes against one another.^45^ With more pressures exerted on these mice (e.g., severe food restriction, limited time budget, an environment with increasing reward scarcity tested across months, dynamic range of offer costs and flavor preferences, as well as a whole host of competing but separately measurable behaviors [offer zone choice conflict, wait zone choice conflict, consumption costs, and travel time]^73^), all mice, even those with a history of stress that might otherwise exhibit an anhedonic phenotype in a simple sucrose preference test, may be pressured to perform well on the Restaurant Row task in which case “anhedonia” may manifest in more subtle ways as opposed to overt underperformance. Indeed, defeated mice were as equally as capable as non-defeated mice in their ability to establish equally ranked flavor preferences and engage in goal-oriented planning behavior in the offer zone. Where defeated most differed from non-defeated mice was during wait zone quit decisions.

Change-of-mind decisions in the wait zone capture a unique situation in which mice have the opportunity to abandon an ongoing investment.^74^ Well-trained mice generally make economically advantageous decisions in the offer zone, appropriately accepting low cost offers and rejecting high cost offers as a function of offer length cued by tone pitch and interacting with subjective flavor preferences. Nonetheless, mice sometimes violate their own decision policies with some degree of noise. These events, accepting expensive offers or rejecting inexpensive offers (decisions one typically would not make), can be viewed as economic mistakes on this task. Previously published reports on the Restaurant Row task found that mice were more likely to accept expensive offers in higher preferred restaurants and that passes made through the offer zone to the wait zone when doing so were quicker and more ballistic.^47^ Thus, the majority of quit decisions in the wait zone that follow these initially economically disadvantageous offer zone enter decisions (or “snap judgments”) can be considered to be corrections of past mistakes once in the wait zone given the opportunity to re-evaluate. Previous studies have demonstrated that these change-of-mind decisions in particular process additional information that may reflect affective components of choice history beyond the value strictly tied to the food reward itself.^47,75^ The first report of Restaurant Row tested rats with electrodes implanted in the orbitofrontal cortex.^46^ Investigators found that following such economic mistakes, neural ensembles in the orbitofrontal cortex represented missed opportunities specifically encoding the previously unselected action.^46^ This is thought to reflect the neural correlate of counterfactual thinking often seen during the experience of regret (processing “the road not traveled”).^76^ Following economic violations, animals appeared to engage in compensatory behaviors to make up for lost time.^46^ Similarly, it was previously demonstrated mice engage in these sort of compensatory behaviors and that over time, mice develop strategies to avoid repeating similar mistakes independent of and separate from reinforcement rate maximization, a process related to regret avoidance observed on the Restaurant Row task.^47^

Appreciating error of one’s own agency is central to understanding some of the affective components of decision-making.^77^ The Restaurant Row task appears to be capable of accessing such cognitive structures. Previous reports of regret-related behaviors on the Restaurant Row task specifically control for this by ensuring animals had access to the appropriate information confirming that their decisions were indeed the suboptimal choices.^37,38,46,47^ When animals were instead informed that alternative unselected options were actually no better than chosen options, representations of regret in the brain were not seen.^46^ Taken together, agency of own’s own actions is a critical component of emotion-cognition interactions that is testable in this neuroeconomic framework.

The sunk cost bias in particular captures unique situations during wait zone quit decisions in which continued investment in an ongoing endeavor is at odds with the more economically advantageous decision to cut losses and move on. Here, we found that mice with a history of social defeat stress displayed an increased sensitivity to sunk costs. If hypersensitivity to sunk costs is in fact dependent to some degree on agency and degree of choice commitment, it may be the case that defeated mice display an increase in the gain of this property of decision ownership and its relationship to loss aversion. It was previously demonstrated that sensitivity to sunk costs in mice, rats, and humans tested on Restaurant Row was only observable after commitment to a choice was made but not while individuals were still engaged in a planning process.^36^ Other reports in the human psychology literature have found that the more an individual’s sense of personal responsibility for a chosen action increases, the more sensitive that person is to sunk costs.^78–80^ Other studies examining internalizing versus externalizing lability factors in humans could explain individual differences in loss aversion based on these personality traits that may be related here to processing of error of one’s own agency during choice conflict.^54^ In depressed individuals, sensitivity to sunk costs have been linked to a resistance to change and in more extreme cases, propensity to attempt suicide.^14,15,81–83^ Based on Leahy’s sunk cost theory of depression, overinvestments in goals thus may be seen as an alternative to the collapse in incentive and motivation that comes with abandoning an ongoing endeavor that depressed individuals struggle with and view as a failure on their part.^14^ Changes in sensitivity to sunk costs in depressed individuals, reflected here in this model, may be related to regret avoidance and the tendency depressed individuals have to confuse past and future costs.^47^

Sunk costs by definition arises from valuing spent resources that cannot be recovered. What might be the neural correlates involved is future-looking versus past-sensitive valuations in the brain? Different behaviors measured on the Restaurant Row task have been linked to separable decision-making systems in dissociable neural circuits.^36,46,47,49,50,52–55,84^ Humans previously tested on the translated version of the Restaurant Row task while being scanned using functional magnetic resonance imaging displayed increased activation of the default mode network specifically when in the offer zone.^55^ Network activity that included prefrontal and hippocampal activation revealed representations of future competing options before a decision in the offer zone was made particularly during instances when sunk costs are being ignored.^36,55^ Similarly, in rats trained on Restaurant Row with electrodes implanted in the prefrontal cortex and hippocampus, activity of neural ensembles that encode spatial information represented sequences of possible future locations animals could choose between when entering the offer zone, again during instances when sunk costs are being ignored, suggesting these neural substrates are involved in future-looking deliberation processes across species.^36,84^ Chemogenetic manipulations using inhibitory designer receptors exclusively activated by designer drugs (DREADDs) expressed in the prelimbic subregion of the prefrontal cortex was able to disrupt hippocampal representations of spatial sequences and planning behaviors in the offer zone.^84^ Less is known about the neural substrates involved during sunk cost-sensitive change of mind decisions in the wait zone. Optogenetic manipulation of infralimbic projections to the nucleus accumbens have been shown to disrupt change-of-mind decisions in the wait zone without affecting offer zone behaviors, suggesting this pathway may be specifically altered in defeated mice displaying hypersensitivity to sunk costs.^52^ In other animal studies, manipulations of the infralimbic cortex has been implicated in augmenting behavioral flexibility and extinction learning, perhaps related to computational properties involved in resistance to change, self-control, or inaction inertia.^85^ In addition to the previous work in orbitofrontal cortex and regret on this task mentioned above, other investigators have found that activity in the amygdala in humans and rodents as well as orbitofrontal projections to the amygdala may be related specifically to loss aversion and effort-related properties of reward seeking behavior and too may be contributory to perseverative aspects of economically disadvantageous endeavors.^6,46,76,86–92^ Taken together, this novel neuroeconomic framework for studying the neural correlates of complex emotion-cognition interactions across species may provide a fresh perspective on the pathophysiology of depression.

In this study, following a single dose of ketamine, hypersensitivity to sunk costs was renormalized in defeated mice. Ketamine is a relatively new rapid-acting antidepressant treatment option presented to patients with treatment-resistant depression.^40–42^ These patients have typically failed multiple monoamine-based psychopharmacologic trials and are considering second– or third-line interventions. To date, these options include electroconvulsive therapy, transcranial magnetic stimulation, vagal nerve stimulation, ketamine, and ongoing experimental trials with psilocybin or lysergic acid diethylamide before ultimately escalating to invasive deep brain stimulation therapy.^4,93^ Ketamine, an anesthetic and psychotomimetic, is a non-competitive antagonist of the *N*-methyl-*D*-aspartate glutamate receptor.^48^ Current theories for how ketamine exerts its antidepressant effects remain unclear. However, the recent hypotheses postulate that ketamine, in addition to modulating cyclic adenosine monophosphate signaling (which is reduced in patients with depression and restored following treatment with SSRIs) mediated through potentiation of adrenergic receptors, rapidly increases the synthesis of brain-derived neurotrophic factor which can lead to an increase in synaptogenesis and can augment the endocytic machinery involved in modulating glutamatergic synaptic transmission.^42,94–96^ While it is clear the molecular effects of even a single dose of ketamine can be widespread and multifactorial, a focus that continues to be an active area of investigation, recent studies looking at brain-wide electrophysiological changes in mice subjected to CSDS as well as following a single dose of ketamine at the same dose used in the present study have found network specific signatures in the brain that are both augmented by stress and responsive to ketamine, a network that is driven by specific activity patterns between the amygdala and nucleus accumbens.^97^ While the molecular and circuit-level effects of ketamine are continuously being explored, we present evidence here for the ability of ketamine to selectively alter past-sensitive information processing in a novel neuroeconomic paradigm, shedding light on some of its potential therapeutic mechanisms on emotion-cognition interactions in the brain. Interestingly, we also found that ketamine also completely abolished the sunk cost bias in non-defeated mice. This brings up potential implications for its use and warrants further studies on the differences between treatment dosing during a major depressive episode, maintenance dosing for patients in remission to prevent relapse, and even prophylactic dosing for at risk populations for preventative measures. By augmenting the outlook of an individual, particularly one that may be at an increased vulnerability to develop depression, to be more forward-looking in their decision-making tendencies, these data suggest a mechanism by which ketamine may be protective at least with how stress might impact propensity to focus on past losses. Current ongoing efforts are focused on understanding individual differences in susceptibility versus resiliency to depression; how this is directly related to sensitivity to sunk costs remains to be explored.^72^

While the CSDS model of depression is widely used, one limitation is that this model has been well optimized for use in male mice. Achieving the same level of aggression-related psychosocial stress induced by a dominant CD-1 mouse is more difficult in females. Current efforts in the field are exploring ways to create psychosocial stress models of depression that are equally useful in both males and females for exploring important sex-differences that are likely at play in depression and will serve as the basis of future follow up studies.^98^ Examining the effects of other types of stress, including early life stress, on neuroeconomic decision-making can be more readily tested in both male and female mice and will be of importance for characterizing the developmental trajectories of individuals who might be at an increased vulnerability to develop depression later in life.^99^ Additional future directions that warrant continued exploration include characterizing the time course of behavioral effects, both the duration of neuroeconomic changes induced by stress as well as the changes induced by ketamine. Furthermore, testing different doses of ketamine as well as different dose schedules on inducing lasting changes in sunk cost-related decision-making processes remain to be explored and whether or not such changes might be protective to prevent future relapse. These data serve as the foundation for exploratory studies in both rodents and humans alike closely examining the neural correlates of sunk costs in healthy and depressed individuals as well as recording from and manipulating the circuit-specific computations that are involved in past-sensitive valuation processes that may be uniquely pathological in depression.

How decisions are selected can be an elusive process. Neuroeconomics is an emerging field that in part focuses on how the physical limits of the brain give rise to the cognitive biases that govern the way we make decisions. Here, we provide data showing how animal models of depression can be enriched by taking a neuroeconomic approach to study behavior. We examined complex emotion-cognition interactions and discovered that stress augments the balance between how information is processed in a past-sensitive versus forward-looking manner when making economic choices. Hypersensitivity to sunk costs induced by chronic social defeat stress was renormalized following a single dose of ketamine, suggesting one mechanism by which ketamine exerts its therapeutic effects involves accessing specific features of complex decision systems – including the balance between past-sensitive vs. future-looking choices – that are heavily influenced by emotional state. Taken together, our data uncover unique computation-specific etiologies of the effects of stress on complex decision-making that can serve as the basis to better understand the psychological drivers of the neurobiological underpinnings of depression and the psychological mechanisms underlying treatment response.

## Acknowledgments

We thank members of the labs of Eric Nestler and Scott Russo for helpful discussion and technical assistance. Open-source illustrations obtained from SciDraw (www.scidraw.io), credit Federico Claudi.

## Funding

National Institute of Mental Health grant L40MH127601 (BMS)

National Institute of Mental Health supplement grant R01MH051399-31S1 (BMS)

Leon Levy Scholarship in Neuroscience, New York Academy of Sciences (BMS)

Burroughs Wellcome Fund Career Award for Medical Scientists (BMS)

Animal Models for the Social Dimensions of Health and Aging Research Network via NIH/NIA R24 AG065172 (BMS)

Brain & Behavior Research Foundation Young Investigator Award 31140 (RDC)

## Author Contributions

Conceptualization: RDC, BMS

Methodology: RDC, BMS

Investigation: RDC, BMS

Data curation: RDC, BMS

Formal analysis: BMS

Visualization: RDC, BMS

Funding acquisition: RDC, BMS

Supervision: RDC, BMS

Writing – original draft: RDC, BMS

Writing – review & editing: RDC, BMS

## References

1 Chang, L. J. & Sanfey, A. G. Emotion, decision-making and the brain. Adv Health Econ Health Serv Res 20, 31–53 (2008).

2 Caceda, R., Nemeroff, C. B. & Harvey, P. D. Toward an understanding of decision making in severe mental illness. J Neuropsychiatry Clin Neurosci 26, 196–213, doi:10.1176/appi.neuropsych.12110268 (2014).

3 First, M. B. Diagnostic and statistical manual of mental disorders, 5th edition, and clinical utility. J Nerv Ment Dis 201, 727–729, doi:10.1097/NMD.0b013e3182a2168a (2013).

4. NIMH. The National Institute of Mental Health. *<*www.nimh.nih.gov/health/statistics/major-depression*>* (2017).

5 Lee, D. Decision making: from neuroscience to psychiatry. Neuron 78, 233–248, doi:10.1016/j.neuron.2013.04.008 (2013).

6 Coricelli, G., Dolan, R. J. & Sirigu, A. Brain, emotion and decision making: the paradigmatic example of regret. Trends Cogn Sci 11, 258–265, doi:10.1016/j.tics.2007.04.003 (2007).

7 Kahneman, D. A perspective on judgment and choice: mapping bounded rationality. Am Psychol 58, 697–720, doi:10.1037/0003-066X.58.9.697 (2003).

8 Tversky, A. & Kahneman, D. The framing of decisions and the psychology of choice. Science 211, 453–458, doi:10.1126/science.7455683 (1981).

9 Loewenstein, G., Rick, S. & Cohen, J. D. Neuroeconomics. Annu Rev Psychol 59, 647–672, doi:10.1146/annurev.psych.59.103006.093710 (2008).

10 Arkes, H. & Blumer, C. The psychology of sunk cost. Organizational Behavior and Human Decision Processes 35, 124–140 (1985).

11 Magalhaes, P. & White, K. G. The effect of a prior investment on choice: the sunk cost effect. J Exp Psychol Anim Learn Cogn 40, 22–37, doi:10.1037/xan0000007 (2014).

12 Chung, S. H. & Cheng, K. C. How does cognitive dissonance influence the sunk cost effect? Psychol Res Behav Manag 11, 37–45, doi:10.2147/PRBM.S150494 (2018).

13 Dijkstra, K. A. & Hong, Y. Y. The feeling of throwing good money after bad: The role of affective reaction in the sunk-cost fallacy. PLoS One 14, e0209900, doi:10.1371/journal.pone.0209900 (2019).

14 Leahy, R. L. Sunk Costs and Resistance to Change. Journal of Cognitive Psychotherapy 14, 355–371 (2000).

15 Moon, H., Hollenbeck, J. R., Humphrey, S. E. & Maue, B. The Tripartite Model of Neuroticism and the Suppression of Depression and Anxiety Within an Escalation of Commitment Dilemma. Journal of Personality 71, 347–368 (2003).

16 Leahy, R. L. Sunk costs: Backward-looking decisions. The Behavior Therapist 32, 137–139 (2009).

17 Westfall, J. E., Jasper, J. D. & Christman, S. Inaction inertia, the sunk cost effect, and handedness: avoiding the losses of past decisions. Brain Cogn 80, 192–200, doi:10.1016/j.bandc.2012.06.003 (2012).

18 Putten, M. V., Zeelenberg, M. & Dijk, E. V. Dealing with missed opportunities: Action vs. state orientation moderates inaction inertia. Journal of Experimental Social Psychology 45, 808–815 (2009).

19 Folk, J. B. et al. Wise Additions Bridge the Gap between Social Psychology and Clinical Practice: Cognitive-Behavioral Therapy as an Exemplar. J Psychother Integr 2016, doi:10.1037/int0000038 (2016).

20 Nestler, E. J. & Carlezon, W. A., Jr. The mesolimbic dopamine reward circuit in depression. Biol Psychiatry 59, 1151–1159, doi:10.1016/j.biopsych.2005.09.018 (2006).

21 Bale, T. L. et al. The critical importance of basic animal research for neuropsychiatric disorders. Neuropsychopharmacology 44, 1349–1353, doi:10.1038/s41386-019-0405-9 (2019).

22 Nestler, E. J. et al. Neurobiology of depression. Neuron 34, 13–25, doi:10.1016/s0896-6273(02)00653-0 (2002).

23 Nestler, E. J. Epigenetic mechanisms of depression. JAMA Psychiatry 71, 454–456, doi:10.1001/jamapsychiatry.2013.4291 (2014).

24 Duman, R. S., Heninger, G. R. & Nestler, E. J. A molecular and cellular theory of depression. Arch Gen Psychiatry 54, 597–606, doi:10.1001/archpsyc.1997.01830190015002 (1997).

25 Lobo, M. K., Nestler, E. J. & Covington, H. E., 3rd. Potential utility of optogenetics in the study of depression. Biol Psychiatry 71, 1068–1074, doi:10.1016/j.biopsych.2011.12.026 (2012).

26 Kalenscher, T. & van Wingerden, M. Why we should use animals to study economic decision making – a perspective. Front Neurosci 5, 82, doi:10.3389/fnins.2011.00082 (2011).

27 Hakimzada, F. A., Gutnik, L. A., Yoskowitz, N. A. & Patel, V. L. Cognitive neuroeconomics: new solutions to old problems. AMIA Annu Symp Proc, 974 (2005).

28 Braeutigam, S. Neuroeconomics--from neural systems to economic behaviour. Brain Res Bull 67, 355–360, doi:10.1016/j.brainresbull.2005.06.009 (2005).

29 Sanfey, A. G., Loewenstein, G., McClure, S. M. & Cohen, J. D. Neuroeconomics: cross-currents in research on decision-making. Trends Cogn Sci 10, 108–116, doi:10.1016/j.tics.2006.01.009 (2006).

30 Schultz, W. Introduction. Neuroeconomics: the promise and the profit. Philos Trans R Soc Lond B Biol Sci 363, 3767–3769, doi:10.1098/rstb.2008.0153 (2008).

31 Smith, D. V. & Huettel, S. A. Decision neuroscience: neuroeconomics. Wiley Interdiscip Rev Cogn Sci 1, 854–871, doi:10.1002/wcs.73 (2010).

32 Sharp, C., Monterosso, J. & Montague, P. R. Neuroeconomics: a bridge for translational research. Biol Psychiatry 72, 87–92, doi:10.1016/j.biopsych.2012.02.029 (2012).

33 Sweis, B. M. & Nestler, E. J. Pushing the boundaries of behavioral analysis could aid psychiatric drug discovery. PLoS Biol 20, e3001904, doi:10.1371/journal.pbio.3001904 (2022).

34 Arkes, H. R. & Ayton, P. The Sunk Cost and Concorde Effects: Are Humans Less Rational Than Lower Animals? Psychological Bulletin 125, 591–600 (1999).

35 Magalhaes, P. & Geoffrey White, K. The sunk cost effect across species: A review of persistence in a course of action due to prior investment. J Exp Anal Behav 105, 339–361, doi:10.1002/jeab.202 (2016).

36 Sweis, B. M. et al. Sensitivity to “sunk costs” in mice, rats, and humans. Science 361, 178–181, doi:10.1126/science.aar8644 (2018).

37 Durand-de Cuttoli, R., et al. A double-hit of social and economic stress in mice precipitates changes in decision-making strategies. Biol Psychiatry, doi:10.1016/j.biopsych.2023.12.011 (2023).

38 Durand-de Cuttoli, R., et al. Distinct forms of regret linked to resilience versus susceptibility to stress are regulated by region-specific CREB function in mice. Science Advances 8, eadd5579, doi:10.1126/sciadv.add5579 (2022).

39 Durand-de Cuttoli, R. et al. Chronic social stress induces isolated deficits in reward anticipation on a neuroeconomic foraging task. biorxiv, doi:10.1101/2022.01.17.476514 (2022).

40 Krystal, J. H., Abdallah, C. G., Sanacora, G., Charney, D. S. & Duman, R. S. Ketamine: A Paradigm Shift for Depression Research and Treatment. Neuron 101, 774–778, doi:10.1016/j.neuron.2019.02.005 (2019).

41 Murrough, J. W. Ketamine as a novel antidepressant: from synapse to behavior. Clinical Pharmacology & Therapeutics 91, 303–309 (2012).

42 Serafini, G., Howland, R. H., Rovedi, F., Girardi, P. & Amore, M. The role of ketamine in treatment-resistant depression: a systematic review. Curr Neuropharmacol 12, 444–461, doi:10.2174/1570159X12666140619204251 (2014).

43 Krishnan, V. et al. Molecular adaptations underlying susceptibility and resistance to social defeat in brain reward regions. Cell 131, 391–404, doi:10.1016/j.cell.2007.09.018 (2007).

44 Zhao, C. X. et al. The hidden opportunity cost of time effect on intertemporal choice. Front Psychol 6, 311, doi:10.3389/fpsyg.2015.00311 (2015).

45 Stephens, D. W. Decision ecology: foraging and the ecology of animal decision making. Cogn Affect Behav Neurosci 8, 475–484, doi:10.3758/CABN.8.4.475 (2008).

46 Steiner, A. P. & Redish, A. D. Behavioral and neurophysiological correlates of regret in rat decision-making on a neuroeconomic task. Nat Neurosci 17, 995–1002, doi:10.1038/nn.3740 (2014).

47 Sweis, B. M., Thomas, M. J. & Redish, A. D. Mice learn to avoid regret. PLoS Biol 16, e2005853, doi:10.1371/journal.pbio.2005853 (2018).

48 Rosenbaum, S. B., Gupta, V. & Palacios, J. L. in StatPearls (2020).

49 Sweis, B. M., Thomas, M. J. & Redish, A. D. Beyond simple tests of value: measuring addiction as a heterogeneous disease of computation-specific valuation processes. Learn Mem 25, 501–512, doi:10.1101/lm.047795.118 (2018).

50 Sweis, B. M., Redish, A. D. & Thomas, M. J. Prolonged abstinence from cocaine or morphine disrupts separable valuations during decision conflict. Nat Commun 9, 2521, doi:10.1038/s41467-018-04967-2 (2018).

51 Carter, E. C. & Redish, A. D. Rats value time differently on equivalent foraging and delay-discounting tasks. J Exp Psychol Gen 145, 1093–1101, doi:10.1037/xge0000196 (2016).

52 Sweis, B. M., Larson, E. B., Redish, A. D. & Thomas, M. J. Altering gain of the infralimbic-to-accumbens shell circuit alters economically dissociable decision-making algorithms. Proc Natl Acad Sci U S A 115, E6347–E6355, doi:10.1073/pnas.1803084115 (2018).

53 Abram, S. V., Breton, Y. A., Schmidt, B., Redish, A. D. & MacDonald, A. W., 3rd. The Web-Surf Task: A translational model of human decision-making. Cogn Affect Behav Neurosci 16, 37–50, doi:10.3758/s13415-015-0379-y (2016).

54 Abram, S. V., Redish, A. D. & MacDonald, A. W., 3rd. Learning From Loss After Risk: Dissociating Reward Pursuit and Reward Valuation in a Naturalistic Foraging Task. Front Psychiatry 10, 359, doi:10.3389/fpsyt.2019.00359 (2019).

55 Abram, S. V., Hanke, M., Redish, A. D. & MacDonald, A. W., 3rd. Neural signatures underlying deliberation in human foraging decisions. Cogn Affect Behav Neurosci 19, 1492–1508, doi:10.3758/s13415-019-00733-z (2019).

56 Rillich, J. & Stevenson, P. A. Serotonin Mediates Depression of Aggression After Acute and Chronic Social Defeat Stress in a Model Insect. Front Behav Neurosci 12, 233, doi:10.3389/fnbeh.2018.00233 (2018).

57 Krishnan, V. & Nestler, E. J. The molecular neurobiology of depression. Nature 455, 894–902, doi:10.1038/nature07455 (2008).

58 Faraji, J. et al. Evidence for Ancestral Programming of Resilience in a Two-Hit Stress Model. Front Behav Neurosci 11, 89, doi:10.3389/fnbeh.2017.00089 (2017).

59 Pena, C. J. & Nestler, E. J. Progress in Epigenetics of Depression. Prog Mol Biol Transl Sci 157, 41–66, doi:10.1016/bs.pmbts.2017.12.011 (2018).

60 Bagot, R. C. et al. Circuit-wide Transcriptional Profiling Reveals Brain Region-Specific Gene Networks Regulating Depression Susceptibility. Neuron 90, 969–983, doi:10.1016/j.neuron.2016.04.015 (2016).

61 Han, M. H. & Nestler, E. J. Neural Substrates of Depression and Resilience. Neurotherapeutics 14, 677–686, doi:10.1007/s13311-017-0527-x (2017).

62 Berton, O. et al. Essential role of BDNF in the mesolimbic dopamine pathway in social defeat stress. Science 311, 864–868, doi:10.1126/science.1120972 (2006).

63 Cao, J. L. et al. Mesolimbic dopamine neurons in the brain reward circuit mediate susceptibility to social defeat and antidepressant action. J Neurosci 30, 16453–16458, doi:10.1523/JNEUROSCI.3177-10.2010 (2010).

64 Barik, J. et al. Chronic stress triggers social aversion via glucocorticoid receptor in dopaminoceptive neurons. Science 339, 332–335, doi:10.1126/science.1226767 (2013).

65 Vialou, V. et al. Serum response factor promotes resilience to chronic social stress through the induction of DeltaFosB. J Neurosci 30, 14585–14592, doi:10.1523/JNEUROSCI.2496-10.2010 (2010).

66 Vialou, V. et al. DeltaFosB in brain reward circuits mediates resilience to stress and antidepressant responses. Nat Neurosci 13, 745–752, doi:10.1038/nn.2551 (2010).

67 Golden, S. A. et al. Epigenetic regulation of RAC1 induces synaptic remodeling in stress disorders and depression. Nat Med 19, 337–344, doi:10.1038/nm.3090 (2013).

68 Hodes, G. E. et al. Individual differences in the peripheral immune system promote resilience versus susceptibility to social stress. Proc Natl Acad Sci U S A 111, 16136–16141, doi:10.1073/pnas.1415191111 (2014).

69 Menard, C. et al. Social stress induces neurovascular pathology promoting depression. Nat Neurosci 20, 1752–1760, doi:10.1038/s41593-017-0010-3 (2017).

70 Menard, C., Pfau, M. L., Hodes, G. E. & Russo, S. J. Immune and Neuroendocrine Mechanisms of Stress Vulnerability and Resilience. Neuropsychopharmacology 42, 62–80, doi:10.1038/npp.2016.90 (2017).

71 Hodes, G. E. et al. Sex Differences in Nucleus Accumbens Transcriptome Profiles Associated with Susceptibility versus Resilience to Subchronic Variable Stress. J Neurosci 35, 16362–16376, doi:10.1523/JNEUROSCI.1392-15.2015 (2015).

72 Cathomas, F., Murrough, J. W., Nestler, E. J., Han, M. & Russo, S. J. Neurobiology of resilience: interface between mind and body. Biological Psychiatry 86, 410–420 (2019).

73 Nwakama, C. A. et al. Diabetes alters neuroeconomically dissociable forms of mental accounting. bioRxiv, doi: doi.org/10.1101/2024.01.04.574210 (2024).

74 Durand-de Cuttoli, R. et al. Sex differences in change-of-mind neuroeconomic decision-making is modulated by LINC00473 in medial prefrontal cortex. bioRxiv, doi:10.1101/2024.05.08.592609 (2024).

75 Resulaj, A., Kiani, R., Wolpert, D. M. & Shadlen, M. N. Changes of mind in decision-making. Nature 461, 263–266, doi:10.1038/nature08275 (2009).

76 Steiner, A. P. & Redish, A. D. The road not taken: neural correlates of decision making in orbitofrontal cortex. Front Neurosci 6, 131, doi:10.3389/fnins.2012.00131 (2012).

77 Howlett, J. R. & Paulus, M. P. Decision-Making Dysfunctions of Counterfactuals in Depression: Who Might I have Been? Front Psychiatry 4, 143, doi:10.3389/fpsyt.2013.00143 (2013).

78 Simonson, I. & Nye, P. The effect of accountability on susceptibility to decision errors. Organ Behav Hum Decis Process 51, 416–446 (1992).

79 Whyte, G. Escalating commitment in individual and group decision making: A prospect theory approach. Organ Behav Hum Decis Process 54, 430–455 (1993).

80 Staw, B. M. Knee-deep in the big muddy: A study of escalating commitment to a chosen course of action. Organizational Behavior and Human Performance 16 (1976).

81 Takahashi, T. Neuroeconomics of suicide. Neuro Endocrinol Lett 32, 400–404 (2011).

82 Jarmolowicz, D. P., Bickel, W. K., Sofis, M. J., Hatz, L. E. & Mueller, E. T. Sunk costs, psychological symptomology, and help seeking. Springerplus 5, 1699, doi:10.1186/s40064-016-3402-z (2016).

83 Szanto, K. et al. Decision-making competence and attempted suicide. J Clin Psychiatry 76, e1590–1597, doi:10.4088/JCP.15m09778 (2015).

84 Schmidt, B., Duin, A. A. & Redish, A. D. Disrupting the medial prefrontal cortex alters hippocampal sequences during deliberative decision making. J Neurophysiol 121, 1981–2000, doi:10.1152/jn.00793.2018 (2019).

85 Barker, J. M., Taylor, J. R. & Chandler, L. J. A unifying model of the role of the infralimbic cortex in extinction and habits. Learn Mem 21, 441–448, doi:10.1101/lm.035501.114 (2014).

86 Camille, N. et al. The involvement of the orbitofrontal cortex in the experience of regret. Science 304, 1167–1170, doi:10.1126/science.1094550 (2004).

87 Camille, N. et al. Striatal sensitivity to personal responsibility in a regret-based decision-making task. Cogn Affect Behav Neurosci 10, 460–469, doi:10.3758/CABN.10.4.460 (2010).

88 Coricelli, G. et al. Regret and its avoidance: a neuroimaging study of choice behavior. Nat Neurosci 8, 1255–1262, doi:10.1038/nn1514 (2005).

89 Coricelli, G. The potential role of regret in the physician-patient relationship: insights from neuroeconomics. Adv Health Econ Health Serv Res 20, 85–97 (2008).

90 Groman, S. M. et al. Orbitofrontal circuits make distinct contributions to flexible decision-making. Neuron 103, 734–746 (2019).

91 Zeng, J., Zhang, Q., Chen, C., Yu, R. & Gong, Q. An fMRI study on sunk cost effect. Brain Res 1519, 63–70, doi:10.1016/j.brainres.2013.05.001 (2013).

92 Fujino, J. et al. Neural mechanisms and personality correlates of the sunk cost effect. Sci Rep 6, 33171, doi:10.1038/srep33171 (2016).

93 Gelenberg, A. J. A review of the current guidelines for depression treatment. J Clin Psychiatry 71, e15, doi:10.4088/JCP.9078tx1c (2010).

94 Yang, Y., Ju, W., Zhang, H. & Sun, L. Effect of Ketamine on LTP and NMDAR EPSC in Hippocampus of the Chronic Social Defeat Stress Mice Model of Depression. Front Behav Neurosci 12, 229, doi:10.3389/fnbeh.2018.00229 (2018).

95 Cavanagh, S. E., Lam, N. H., Murray, J. D., Hunt, L. T. & Kennerley, S. W. A circuit mechanism for decision-making biases and NMDA receptor hypofunction. Elife 9, doi:10.7554/eLife.53664 (2020).

96 Sleigh, J., Harvey, M., Voss, L. & Denny, B. Ketamine – More mechanisms of action than just NMDA blockade. Trends in Anaesthesia and Critical Care 4, 76–81 (2014).

97 Hultman, R. et al. Brain-wide Electrical Spatiotemporal Dynamics Encode Depression Vulnerability. Cell 173, 166–180 e114, doi:10.1016/j.cell.2018.02.012 (2018).

98 Harris, A. Z. et al. A Novel Method for Chronic Social Defeat Stress in Female Mice. Neuropsychopharmacology 43, 1276–1283, doi:10.1038/npp.2017.259 (2018).

99 Pena, C. J. et al. Early life stress confers lifelong stress susceptibility in mice via ventral tegmental area OTX2. Science 356, 1185–1188, doi:10.1126/science.aan4491 (2017).

